# Genetic population variation and phylogeny of *Sinomenium acutum* (Menispermaceae) in subtropical China through chloroplast marker

**DOI:** 10.1101/449900

**Authors:** Ying He, Chun Guo, Xiyao Zeng, Hua Yang, Xingyao Xiong, Ping Qiu

**Author notes:** Corresponding author (Hua Yang), (Hua Yang). These authors contributed equally to this work (Ying He, Chun Guo).

## Abstract

*Sinomenium acutum* (Menispermaceae) is a traditional Chinese medicine. In recent years, extensive harvesting for medicinal purposes has resulted in a sharp decline in its population. Genetic information is crucial for the proper exploitation and conservation of *Sinomenium acutum*, but little is known about it at present. In this study, we analyzed 77 samples from 4 populations using four non-coding regions (*atp*I-*atp*H, *trn*Q-5’*rps*16, *trn*H-*psb*A, and *trn*L-*trn*F) of chloroplast DNA and 14 haplotypes (from C1 to C14) were identified. C1 and C3 were common haplotypes, which were shared by all populations, and C3 was an ancestral haplotype, the rest were rare haplotypes. Obvious phylogeographic structure was not existed inferred by *G*_ST_ / *N*_ST_ test. Mismatch distribution, Tajima’s D and Fu’s F_S_ tests failed to support a rapid demographic expansion in *Sinomenium acutum*. AMOVA highlighted that the high level of genetic differentiation within population. Low genetic variation among populations illustrated gene flow was not restricted. Genetic diversity analyses demonstrated that the populations of Xuefeng, Dalou, and Daba Mountains were possible refugia localities of *Sinomenium acutum*. Based on this study, we proposed a preliminary protection strategy for it that C1, C3, C11 and C12 must be collected. These results offer an valuable and useful information for this species of population genetic study as well as further conservation.

## Introduction

*Sinomenium acutum*, also known as ‘qingteng’, ‘xunfengteng’, and ‘dianfangji’, etc, belongs to the genus *Sinomenium* and the family Menispermaceae. It includes *Sinomenium acutum* (Thunb.) Rehd. et Wils. and *S. acutum* (Thunb.) Rehd. et Wils. var. *cinereum* (Diels) Rehd. et Wils., as recorded in the Pharmacopoeia of the People’s Republic of China (PPRC). *S. acutum* is a traditional Chinese medicine that has been widely used to treat arthromyodynia, rheumatism, and similar diseases for more than one thousand years. It has been reported to have anti-inflammatory, anti-arrhythmic, depression, anti-rheumatic, anti-angiogenic, anti-anxiety, immunosuppressive, antihypertensive, and vasodilating effects (Nishida and Satoh. 2007; Ou et al. 2011; Song et al. 2010; Wang and Li. 2011; Yi et al. 2012).

*S. acutum* grows over a vast geographical area in China (Zhao et al. 2005). In the early stage, we determined the specific distribution range and situation of it by consulting relevant data and field trips. In China Mainland, its roughly horizontal distribution ranges are from 103°76’E to 121°75’ E, and from 23°24’N to 34°51’N. Distribution areas are mainly mountainous, such as Qinling Mountains (QL), Dabie Mountains, Daba Mountains (DB), Wushan Mountains, Dalou Mountains (DL), Luoxiao Mountains (LX), Xuefeng Mountains (XF), Wuyi Mountains, Nanling Mountains (Fig. 1). The climate type is mainly subtropical monsoon in this region. The region has a mild climate, complex terrain, extremely rich in ancient plant lineages (Axelrod et al. 1996), and high species diversity (Myers et al. 2000; Qian and Ricklefs. 2000). The varied terrain of subtropical China provides different microhabitats for living organisms, which is considered as one of the most important refugia for these ancient lineages in the middle Miocene (about 15 Ma) (Axelrod et al. 1996), and it is also the main reason for the high biodiversity compared to other parts of this region (Wu. 1980; Ying et al. 1993; Ying. 2000).

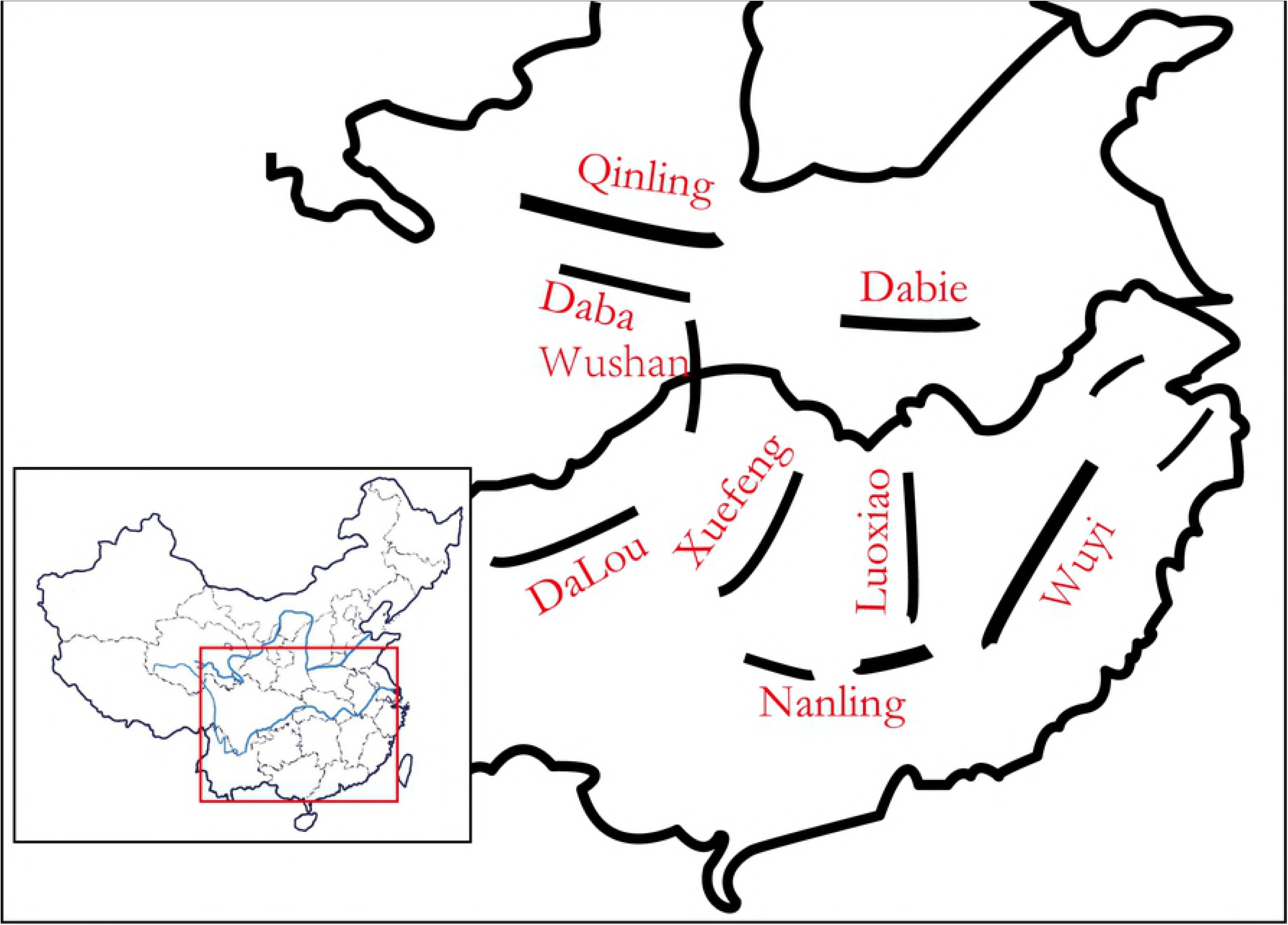

*S. acutum* has strong adaptability to the environment, but high demand for it has led to uncontrolled exploitation in recent years, causing a sharp population decline. It is therefore urgent to establish appropriate measures of exploitation and protection for *S. acutum*. Phylogeography has become one of the main research methods for years (Liu et al. 2012; Qiu et al. 2011) used to better understand the evolutionary history of Chinese subtropical flora and provide a basis for protection policy and management. Studies on *S. acutum* were mainly focused on its pharmacological effects, while its genetic population variation and phylogeny have not yet been conducted except for one of our recent studies based on nuclear ribosomal internal transcribed spacer 2 marker. Phylogeographic studies of *S. acutum* should be carried out in order to better understand the gene genealogies, population variation, and lay the foundation for the formulation of effective conservation strategies.

In this research, we apply cpDNA as a molecular marker for phylogenetic analysis, because the cpDNA has the following characteristics, for instance slow evolution, non-recombinant, maternally inherited, and dispersed via seed (Corriveau and Coleman. 1988; Wolfe et al. 1987).

## Materials and methods

### Sampling

Leaf samples were collected from five populations. Detailed informations, such as population location, number of samples, latitude, and longitude, were listed. Samples were dried in silica gel in the field, and then stored in an ultracold refrigerator (™80°C) until DNA extraction.

### DNA extraction and PCR amplification

The genomic DNA of *S. acutum* was extracted using Plant Genomic DNA Extraction Kit (Tiangen Biotech (Beijing) Co., Ltd.). The concentration and purity of extracted total DNA were determined using a bioanalyzer and 1.0% agarose gel, and then the DNA was diluted with deionized-distilled water and stored at 4°C.

We carried out PCR amplification for four intergenic spacers of cpDNA (*atp*I-*atp*H, *trn*Q-5’*rps*16, *trn*H-*psb*A, as well as *trn*L-*trn*F) (Cieslak et al. 2005; Shaw et al. 2005, 2007). The primer sequences were listed in Table 1. PCR amplification was performed using 25-μL reaction system, containing 40 ng DNA template, 2.5 μL 10 × PCR buffer, 1.0 U Taq DNA polymerase, 2.5 mM Mg^2+^, 0.3 mM dNTPs, and 0.2 μM of each primer. PCR reaction condition was as follows: 4 minutes at 95°C; 35 cycles of 30 seconds at 94°C, 1 minute at 55°C, and 1 minute at 72°C; followed by final extension for 10 minutes at 72°C. The PCR amplification product was assessed by 1.5% agarose gel electrophoresis in 1× TAE buffer.

**Table 1.**
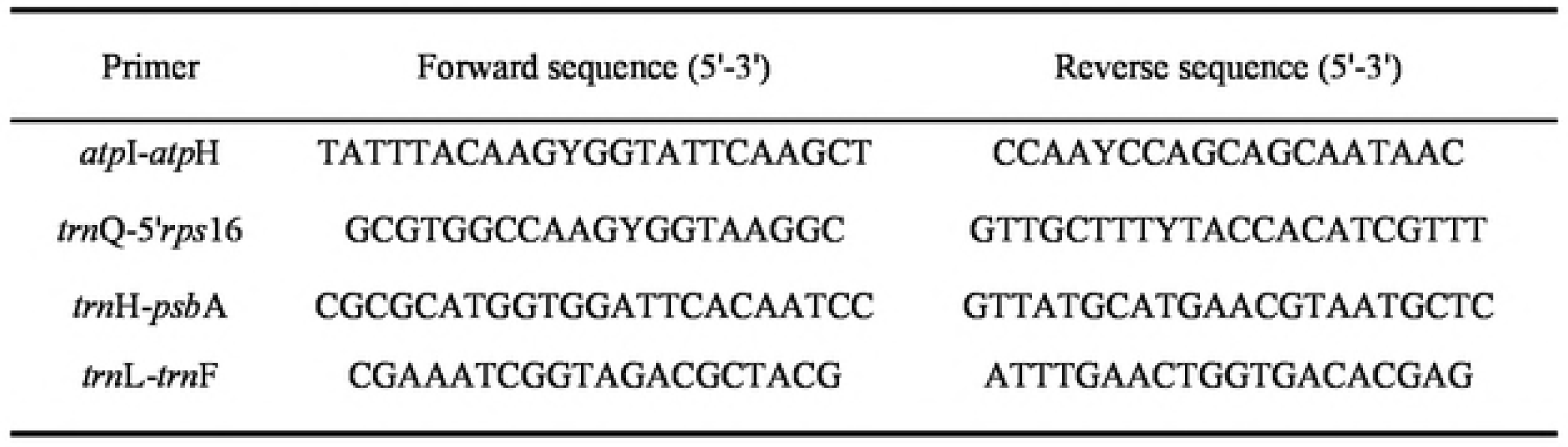
The primer sequences of noncoding chloroplast DNA regions

### Analysis of sequence data

All sequences were aligned using Mega version 6.06 (Tamura et al. 2013), and the misread bases were adjusted manually before aligning using Chromas. Polymorphic sites, number of haplotypes, haplotype diversity (*H*d), and nucleotide diversity (π) were accomplished in DnaSP v5.0 (Librado and Rozas. 2009). A recent demographic history *of S. acutum* was investigated using mismatch distribution analysis in DnaSP v5.0 to assess whether intraspecific lineages experienced past population expansions.

Total genetic diversity (*H_T_*), genetic diversity in within-population (*H*_S_), interpopulation differentiation (*G*_ST_: based only on haplotype frequency), and number of substitution types (*N*_ST_: based on both haplotype frequency and genetic mutation steps among haplotypes) were calculated using the program PERMUT v2.0 (Pons and Petit. 1996). If the value of *N_ST_* was significantly larger than *G*_ST_, we concluded that an obvious phylogeographic structure was present, which means that closely related haplotypes are found more often in the same area than distantly related ones. The proportions of total genetic variance among and within populations were calculated by the analysis of molecular variance (AMOVA) framework in ARLEQUIN v3.5.2.2 (Excoffier and Lischer. 2010; Excoffier et al. 1992), and the significance was tested by 1000 permutations (Sarma et al. 2012; Shepherd et al. 2016; Yu et al. 2014). In addition, Tajima’s D and Fu’s F_S_ tests were performed to evaluate the hypothesis of population expansion. Gene flow (*N*m) was estimated using the equation *N*m = (1 − *F_ST_*) / 2*F_ST_*.

A haplotype network was constructed using the Median Joining model in TCS1.21 (Clement et al. 2000) to delineate intraspecific relationships of cpDNA haplotypes.

### Data available

https://pan.baidu.com/s/1nvHxaiP password: 2nth

## Results

### DNA polymorphisms and haplotype diversity

Four cpDNA fragments *atp*I-*atp*H, *trn*Q-5’*rps*16, *trn*H-*psb*A, and *trn*L-*trn*F were sequenced from 77 individuals of *S. acutum.* The combined length of the four segments ranged from 3658 to 3681 bp, with a consensus alignment length of 3692 bp. Fourteen haplotypes were identified from all *S. acutum* individuals (Fig. 2) based on 12 polymorphism sites (regions) (Table 2), which included 6 nucleotide substitutions and 6 indels. All populations, 3 populations with 5 haplotypes, and 1 population with 6 haplotypes. Haplotype C1 and C3 were the predominant, shared by all populations. Eleven of the haplotypes were endemic (Fig. 2, Table. 3). The total nucleotide diversity (π) and haplotype diversity (*Hd*) in all populations were 2.19 × 10-3 and 0.7553, respectively. Population of XF had the highest nucleotide diversity and haplotype diversity (3.25 × 10-3, 0.9048, respectively), followed by DL and DB (1.75 × 10-3, 0.8312, and 2.87 × 10-3, 0.6429, respectively) (Table 3). These results revealed that the level of genetic differentiation was high in among populations. The geographical distribution of cpDNA haplotypes is shown in Fig. 2.

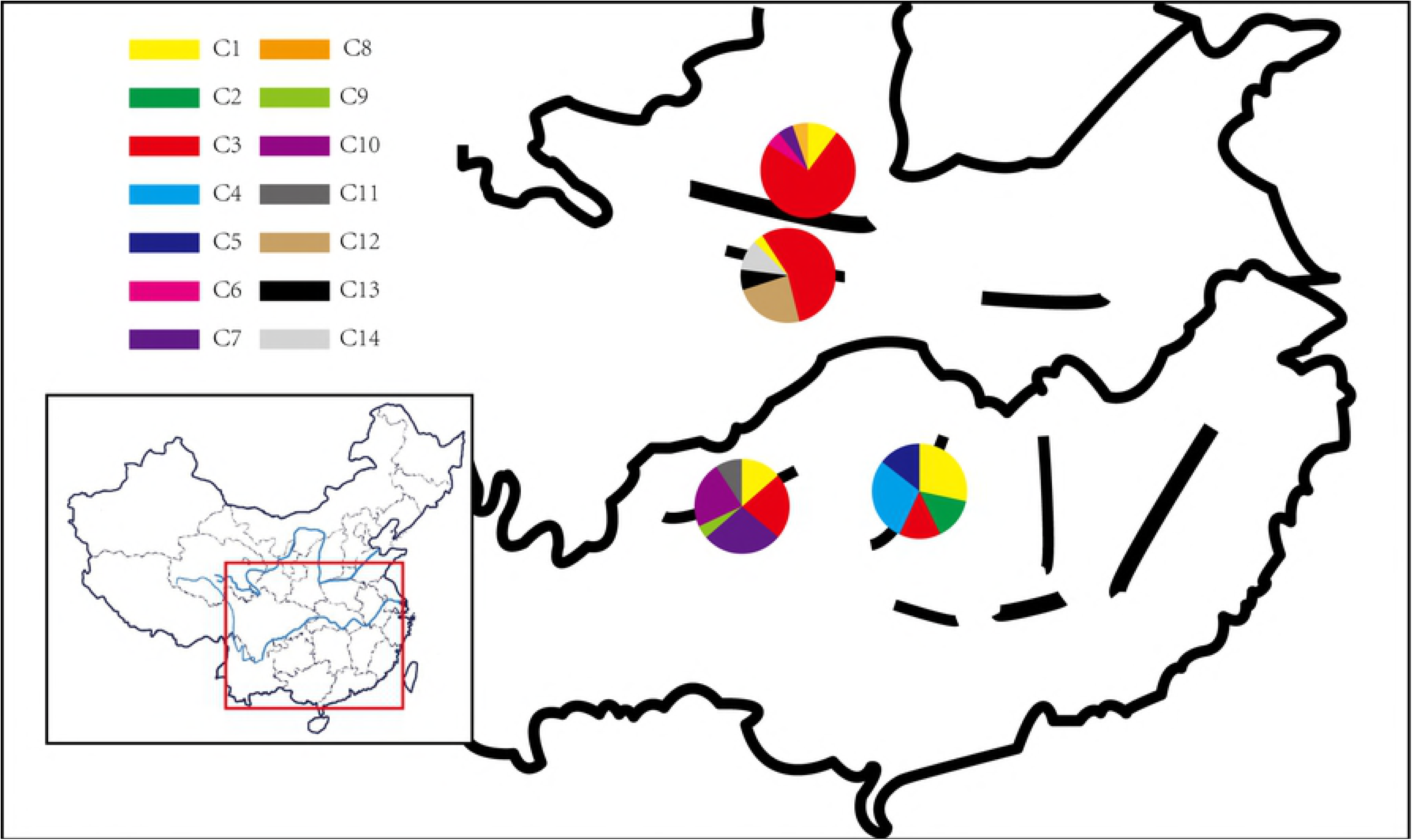

**Table 2.**
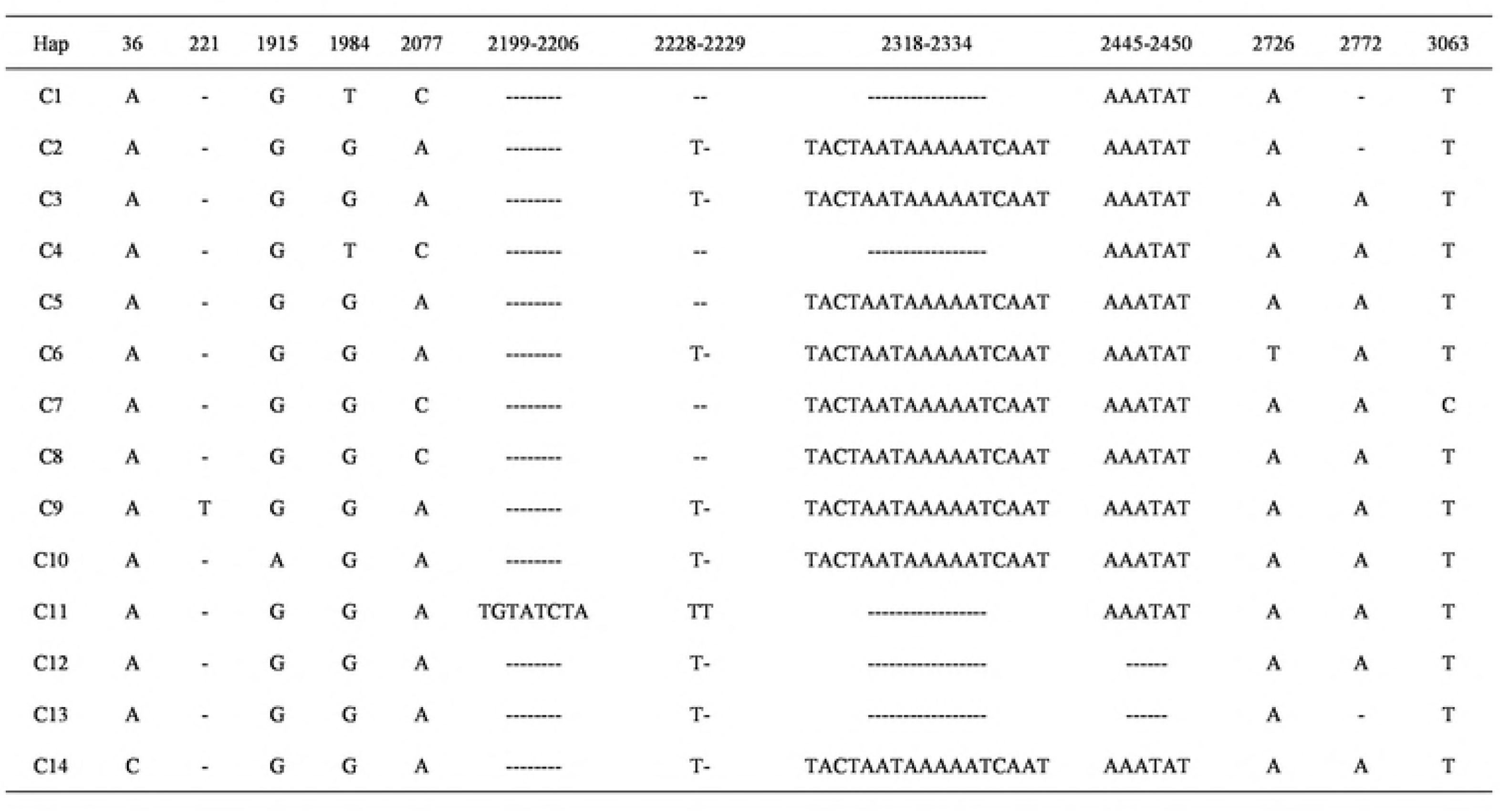
Variable nucleotide siles in 14 haplotypcs of *Smometuum aculum*

**Table 3.** Sample information and haplolypc distribution of *Sinomenium acutum*

| Population (code) | Longitude (°E) | Latitude (°N) | Sample size | Haplotypes (No. of individuals) | *Hd* | $\pi/10^{-3}$ |
| --- | --- | --- | --- | --- | --- | --- |
| Qinling Mountains (QL) | 107.24 | 34.36 | 19 | C1(2), C3(14), C6(1), C7(1), C8(1) | 0.4620 | 0.22 |
| Daba Mountains (DB) | 112.12 | 32 | 29 | C1(1), C3(16), C12(7), C13(2), C14(3) | 0.6429 | 2.87 |
| Xuefeng Mountains (XF) | 110 | 27.57 | 7 | C1(2), C2(1), C3(1), C4(2), C5(1) | 0.9048 | 3.25 |
| Dalou Mountains (DL) | 106.93 | 27.73 | 22 | C1(3), C3(5), C7(6), C9(1), C10(5), C11(2) | 0.8312 | 1.75 |

### Population genetic structure

*H*_T_ was greater than *H*_S_ (*H*_T_ = 0.828, *H*_S_ = 0.710), revealed the high level of genetic differentiation in all populations. A permutation test illustrated that there was no significant difference between *G*_ST_ and *N*_ST_ (*N*_ST_ = 0.139, *G*_ST_ = 0.142; *P* > 0.05). And the value of *N*_ST_ was not significantly larger than *G*_ST,_ therefore, the null hypothesis of a strong phylogeographic pattern was rejected, which showed that haplotypes had little correlation with the geographical areas (Table 4).

**Table 4.** Estimates of wilhin population diversity (*H*_s_), total gene diversity (*H*T)> intcrpopulation genetic difTcrcnlialion (Gyp), number of subslilulion types (Afar), and mismatch distribution analysis

| Population | $H_T$ (se) | $H_S$ (se) | $G_{ST}$ | $N_{ST}$ | P | Tajima's D test | | Fu's Fs test | |
| --- | --- | --- | --- | --- | --- | --- | --- | --- | --- |
|  |  |  |  |  |  | D | p | Fs | p |
| All | 0.828<br>(0.0913) | 0.710<br>(0.0994) | 0.142 | 0.139 | 0.441<br>NS | -0.97017 | P > 0.1<br>NS | 3.32772 | P > 0.1<br>NS |

NS, not significant

AMOVA revealed that 16.38% of the total variation was distributed at among populations (*F*_ST_ = 0.16383), 83.62% existed in within populations (Table 5). These results showed that high genetic differentiations existed in within population. The value of *N*m is 2.552, indicating that the level of gene exchange in among populations was extremely frequent.

**Table 5.** Analysis of molecular variance (AMOVA) for the 4 populations of *Sinomenium aculum*

| Source of variation | *d.f.* | Sum of squares | Variance components | Percentage of variation |
| --- | --- | --- | --- | --- |
| Among populations | 3 | 48.540 | 0.69563Va | 16.38 |
| Within populations | 73 | 259.174 | 3.55033Vb | 83.62 |
| Total | 76 | 307.714 | 4.24596 |  |
| $F_{ST} = 0.16383$ | | | | |

### Phylogenetic relationships among haplotypes

The haplotype network of *S. acutum* was defined (Fig. 3). Haplotype C3 was in the central position of network and was the most widespread haplotype, thus, it may be an ancestral haplotype for all populations. The eleven rare haplotypes (C2, C4, C5, C6, and from C8 to C14) were identified in only a single population, C7 was appeared in two populations. The twelve haplotypes can be obtained by one or multiple mutational step from C3, therefore it making a “star-like” relational network of haplotypes (Fig. 3). The network did not reveal any significant geographical patterns, for instance, haplotype H12, H13, and H14 were distributed in same geographical population, however, H14 has a large genetic distance with H12 and H13, it means the closely related haplotypes are not found more often in the same area than distantly related ones at all.

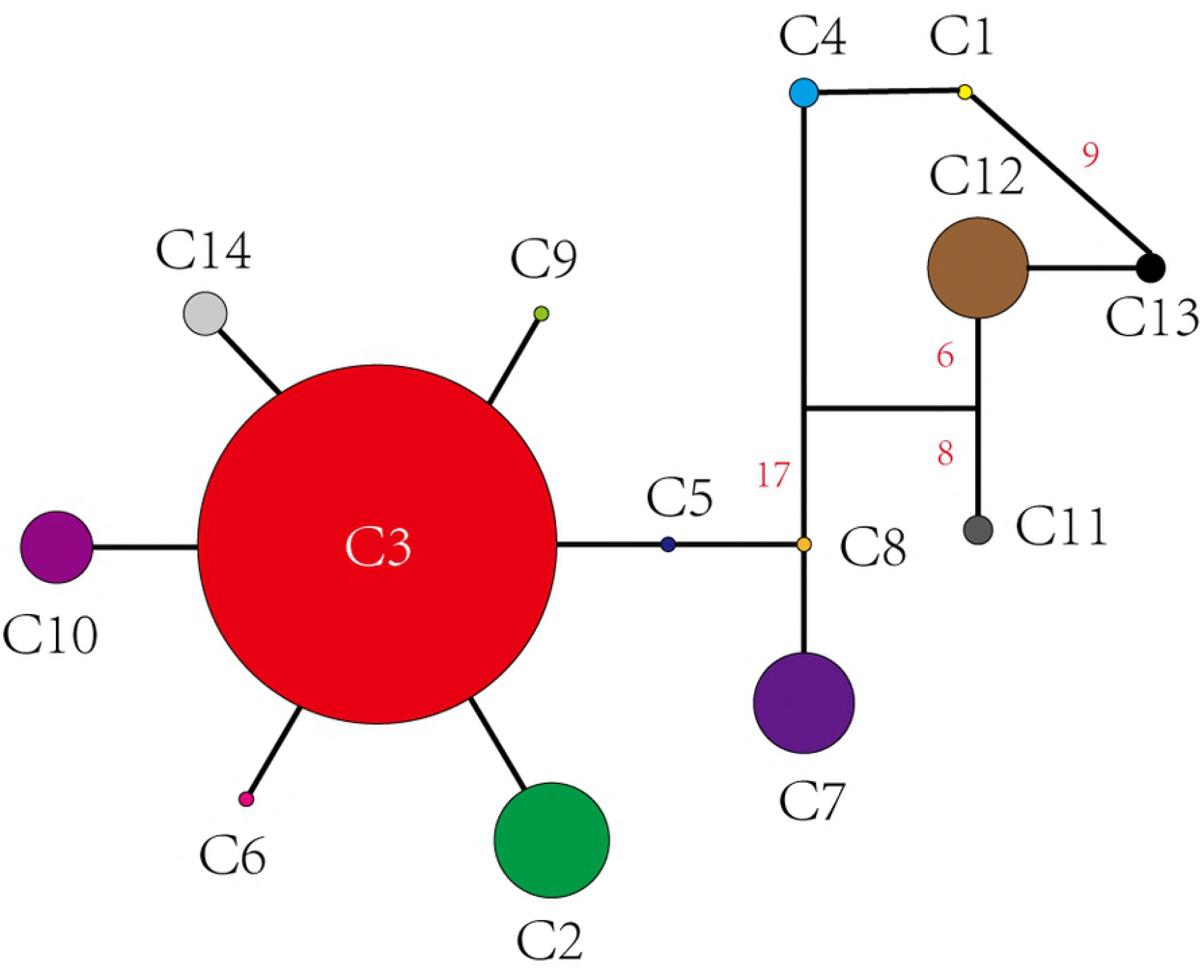

### Demographic history analyses

The mismatch distribution of pairwise nucleotide differences exhibited that the observed curve was inconsistent with the expected curve for all populations, and exhibited a multimodal (Fig. 4). The values of Tajima’s D (D = − 0.97017, *P* > 0.1) and Fu’s F_S_ (Fs = 3.32772, *P* > 0.1) were not significantly for all populations (Table 4). These results provide evidence that *S. acutum* did not experience a recent population expansion.

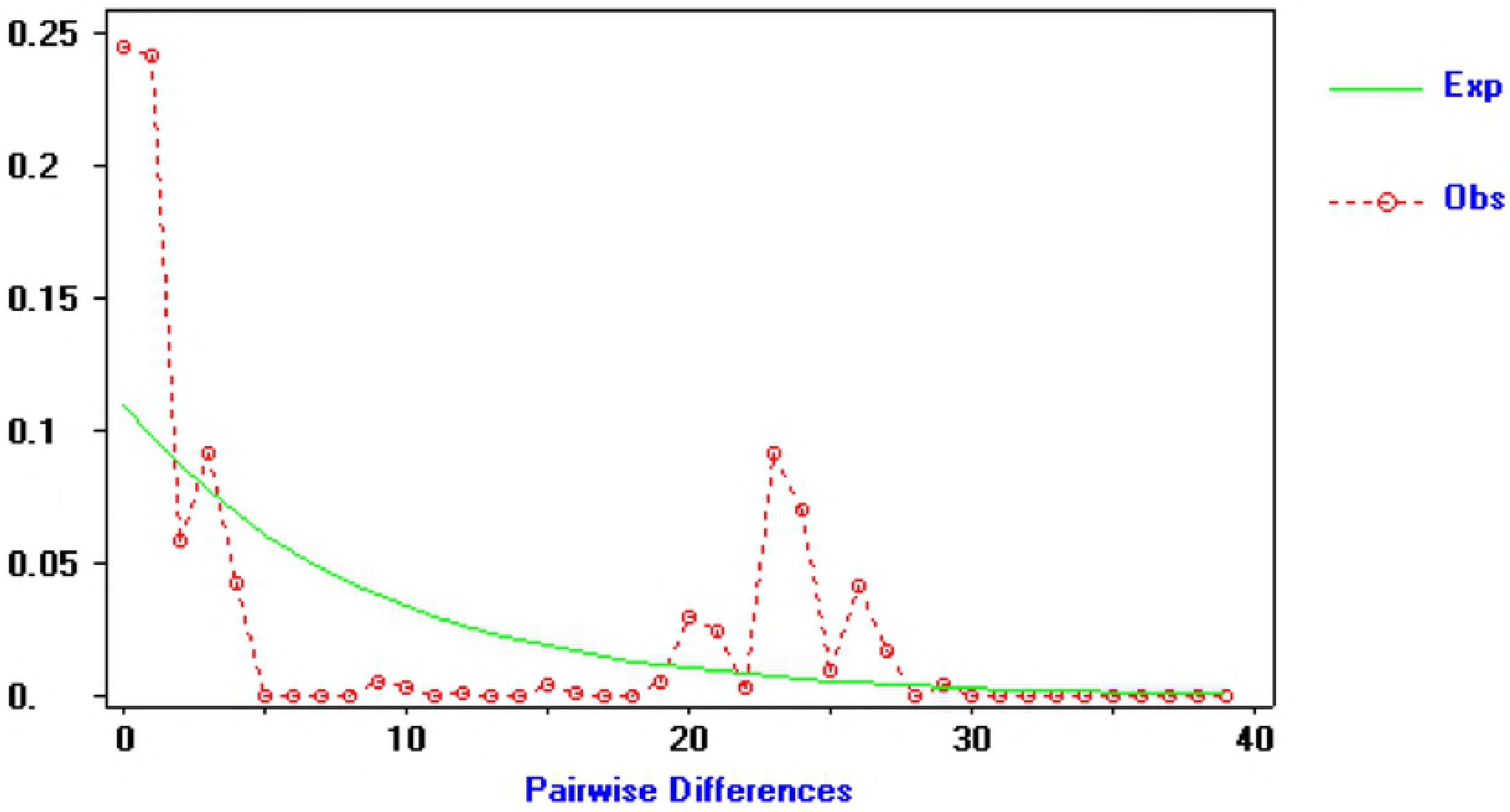

## Discussion

### Phylogenetic relationships of haplotypes

Haplotype C1 and C3 were common, shared by all populations. Since C3 was positioned at the center of the haplotype network and identified in all populations, it was inferred to be ancestral haplotype. Conversely, the twelve haplotypes were surrounded the C3. Moreover, different mutation steps occurred between the ancestral and derived haplotypes. Six haplotypes (C2, C5, C6, C9, C10 and C14) had one mutational step from C3, haplotype C8 had two mutational steps from C3, and C7 had three mutational steps from C3, C1 had 19 mutational steps from it, and C4 had 18 mutational steps, the haplotypes of C11, C12, C13 exhibited larger numbers of mutational steps, indicating that derivation from the ancestral state does require various time.

According to the haplotype network diagram, the haplotypes are in a radial distribution, that is, the other haplotypes are distributed in a divergent manner around the central haplotype with shorter branch lengths. Combining with the distribution of haplotypes, we can conclude that the population has undergone a barely obvious small-scale regional expansion under the appropriate conditions. However, due to the limited ability of population diffusion and the lack of sufficient time after population expansion to established more complex genetic structure, yielded conflicting results. Therefore, we think the most accurately explained that after the glacial period, the population did not experience a significant rapid expansion under optimum conditions.

### Genetic diversity and differentiation

These results, the low genetic variation among populations and high gene flow in the distribution regions, are consistent with these obtained by our previous use of the nuclear ribosomal internal transcribed spacer 2 (ITS2) sequence, but the degree of differentiation was larger than that from ITS2. We suspect that this may due to the genetic characteristics of cpDNA. First, cpDNA is non-recombinant, maternally inherited, and transmitted only through seed, resulting in a slower rate of mutation (Corriveau and Coleman. 1988; Wolfe et al. 1987). In contrast, since nuclear ribosomal DNA (nrDNA) is inherited by parents, the genetic materials is dispersed via seeds and pollen (Hare. 2001), making the mutation rate faster. Thereby, the cpDNA showed greater genetic differentiation than the nuclear DNA. Second, the low ability of seed propagation is the key for the high chloroplast divergence detected in *S. acutum*. Even if birds are fond of eating its fruit would speed up seed dispersal, the high mountains, gorges, and complex paleodrainage basins may be effective barriers to the dispersal and genetic diversification of organisms, which has been reported in multiple phylogeographic studies (He and Jiang. 2014; Yue et al. 2012; Zhang et al. 2010). Third, when cpDNA is used to research gene flow, the effects of male spread are often overlooked.

Neither in the haplotype distribution map nor in the *N*_ST_ / *G*_ST_ test, we failed to detect *S. acutum* with a clear phylogeographic structure. A large number of studies in mainland China show that differential haplotypes usually emerge well subgroups by geographic distance (Liu et al. 2006; Wang et al. 2009). However, we did not found an obvious phylogeographic structure in this study.

Bimodal or multimodal mismatch distributions of pairwise differences between individuals indicate that the population is in a state of equilibrium with a relatively stable size over time, whereas unimodal distributions suggest that the population has recently experienced demographic expansion (Harpending. 1994; Slatkin and Hudson. 1991; Xu et al. 2009; Xue et al. 2014). Significant and non-zero values in Tajima’s D and Fu’s Fs tests may indicate population expansion (An et al. 2015; Fu. 1997; Jin et al. 2016; Kizawa and Maki. 2016; Tajima. 1989; Wen et al. 2015; Xu et al. 2015). Multimodal mismatch distribution curve, Tajima’s D and Fu’s F_S_ values failed to support a recent demographic expansion in *S. acutum*, which are consistent with the results that we obtained from ITS2 sequence.

### Glacial refugia analyses

Glacial refugia is not only the place where animals and plants escape the harsh climate during the glacial period, but also the starting point of species redistribution after glacial period. Regions with high genetic diversity are likely to be the shelters of the glacial age, because these populations usually experience longer population dynamics history than those after glacial expansion. The populations of XF, DL, and DB have rich genetic diversity may reflect these were possible refugia localities. However, we believe that more case studies are needed to increase our knowledge about the phylogeographic history of *S. acutum* in all distribution regions by sufficient sampling.

### Develop conservation measure

The nrDNA reflects both the seed and pollen flow, while the cpDNA only reflects the seed flow. The AMOVA result of cpDNA shows that the seed flow is 2.552 and the seed flow is far less than the pollen flow. Hence, we should mainly rely on the genetic distance of cpDNA haplotype, and combined with the distribution characteristics of haplotype to develop the protection strategy of *S. acutum*. C1 and C3 are common haplotypes, C3 is an ancient haplotype that must be collected, and the genetic relationship of C1 is far apart with C3, thereby, it is feasible to collect its main distribution area. Haplotypes C2, C5, C6, C7, C8, C9, C10, and C14 are close to C3, so these can be considered not saved; C11 and C12 are far away from other haplotypes, consequently, need to hold them; C4 has a close genetic relationship with C1, and C13 is near C12, due to C1 and C12 have been preserved, C4 and C14 are not need to collected.

At present, unfortunately, there are no data to support these views, as study on its genetic diversity and population structure have not yet been conducted. *S. acutum* was distributed in vast geographic areas of China, but we only collected few populations and some populations with little samples due to several problems, which might result in inaccurate results. To better understand the population variation and phylogeny conditions of *S. acutum*, it is necessary to conduct further phylogeographic study based on a large number of nuclear DNA variation to compensate for the drawbacks that cannot be illustrated by chloroplast DNA.

## Conclusion

In order to better understand the genetic diversity and population variation of *S. acutum*, and establish a basis for developing effective conservation strategies, we sequenced four non-coding regions of cpDNA and identified 14 haplotypes from 77 individuals sampled from 4 locations. C1 and C3 were mainly haplotypes, and C3 was inferred to be ancestral haplotype. C7 was occurred only in two populations, and the rest were unique. In addition, we have illustrated the genetic differentiation was occurred mainly in within population, and gene flow was frequent among populations. At last, our study supported that the population was no distinct geographical structure, failed to undergo an obvious rapid expansion, and Xuefeng, Dalou, Daba Mountains seems to be refuges.

## Acknowledgements

The authors would like to thank Jinjing Teng, Tao Guo, Yuan Chen, Shuaifu Li (College of Bioscience and Biotechnology, Hunan Agricultural University) for their help in experiment. We are especially thankful to three anonymous reviewers for their constructive comments which undoubtedly improved this manuscript.

This study was supported by the Hunan Provincial Major Science and Technology Project (S2015S501P010), Hunan Provincial Science and Technology Plan (2015RS4059).

